# Positioning Personal Polygenic Risk score against the population background

**DOI:** 10.1101/813170

**Authors:** Ganna Leonenko, Emily Baker, Karl Michael Schmidt, Valentina Escott-Price, the Alzheimer’s Disease Neuroimaging Initiative

## Abstract

The polygenic risk scores (PRS) approach has been widely used across different traits for estimating polygenic risk, pleiotropy and disease prediction, but mostly in European populations. The predictive ability of the PRS in non-European populations is currently limited due to the lack of genetic research performed in populations of non-European ancestry. One of the main challenges of the practical use of PRS is to place an individual’s personal score in the context of the PRS distribution in the underlying population. In this paper we present an approach for estimating the parameters of the PRS distribution in a population using summary information from public data.

Unstandardized PRS are usually not directly comparable even between European studies. Our approach can be used for standardisation whilst accounting for genotyping platforms, data quality and ancestry. It can be applied to assessing polygenic disease risk for individuals from a European population for any complex genetic disorder and, assuming that most of the disease risk loci are likely to be shared between populations, to estimating the disease risk for individuals from other populations. We demonstrate the precision of our method with simulations. We show the utility of our estimates in application to Alzheimer’s disease in the Alzheimer’s Disease Neuroimaging Initiative (ADNI) study. We present population specific PRSs for different populations using 1000 Genomes data.

## Introduction

Polygenic Risk Score (PRS) has shown potential to stratify individuals into risk categories based on their genetic profile for common genetic disorders. The utility of risk scores in precision medicine is an open question. Application of PRSs in Alzheimer’s Disease (AD) have shown up to 84% prediction accuracy and opened new perspectives for clinical trials. To estimate the disease risk of a particular individual from a particular population, their PRS has to be aligned with a matching background sample where individual genotypes for each SNP are available for each person. The obtained PRS is normally distributed and its mean and variance depend on the number of SNPs and their characteristics (allele frequencies and linkage disequilibrium structure in the population).

The PRS approach has been widely used across different traits, but mostly in European populations. Originally PRSs were designed to summarise genome-wide genotype data into a single variable that measures genetic liability to a disorder or a trait^1^. PRS studies often reach quite high statistical significance levels (small p-value) to suggest trait polygenicity, but the prediction accuracy is usually not sufficient for clinical utility^2^. For example, the prediction accuracy from PRS in schizophrenia has an Area Under the Curve (AUC) of about 70%^3^ and in bipolar disorder an AUC~65%^4^. Nevertheless, PRS was suggested to be a useful tool for the selection of individuals for clinical trials in individuals of European ancestry across different traits^5–8^.

The PRS prediction accuracy of risk for Alzheimer’s disease (AD) is relatively high, especially if the diagnosis is based upon pathology confirmed (AUC is up to 84%) rather than clinical assessment^9^. Furthermore, PRS improves the prediction accuracy significantly over and above *APOE* (which is the strongest predictor of late onset AD risk) and genome-wide significant SNPs^10^. To dissect the disease pathology, the PRS is now being frequently studied for biologically relevant pathways^11^, and can potentially be advantageous for translational studies, e.g. prioritizing samples for stem cell research.

The predictive ability of the PRS in non-European populations is currently limited due to the lack of genetic research performed in populations of non-European ancestry. Previous studies have attempted to assess PRS accuracy across different populations using well-powered European Genome-Wide Association Study (GWAS) discovery cohorts, but have shown mixed results. Reisberg *et al.* compared the distribution of PRSs for type 2 diabetes (T2D) and coronary heart disease (CHD) for different populations and reported lowest scores for the European population compared to Africans^12^. They concluded that PRS models using effects from European-centred GWAS cannot be directly applied to different ancestries for disease prediction. Similarly, Martin *et al.*, by applying PRS from single-ancestry GWAS to eight phenotypes in diverse populations, reported directional differences in all scores and the strongest correlation of true and observed risk in the population from which the effect sizes were derived^13^. It is likely that the main contributors to these differences are the SNP allele frequencies and linkage disequilibrium patterns which differ substantially across populations. The disease association effect sizes are also likely to be different, however, the risk loci are often comparable across populations. For example, Li et al (2017) have shown that about 95% of the genome-wide significant index alleles (or their proxies) from the Psychiatric Genetic Consortium (PGC2) study of European ancestry were overrepresented in Chinese schizophrenia cases, including ∼50% that achieved nominal significance^14^. The Population Architecture using Genomics and Epidemiology (PAGE) study presented consistent association in the same direction for all 19 T2D variants discovered in European cohorts across different ethnic groups^15^. In a more recent PAGE publication^16^, genome-wide association analyses were performed on 49,839 individuals with different ancestry background across 26 traits. They replicated 574 GWAS variant-trait associations in 261 distinct regions of which 132 had significant evidence of effect heterogeneity by genetic ancestry. Therefore gene discoveries and improvement of the disease prediction accuracy for clinical trials still highly rely on powerful GWAS studies in diverse populations.

For dementia related phenotypes, Marden et al (2014) tested whether AD PRS can predict dementia and memory function in non-Hispanic black and white participants^17^ and has shown that the PRS (which includes *APOE*) is elevated in dementia cases, with direction of association being similar for both samples.

It has also been suggested that causal effect estimates tend to be shared between populations^18^; however the prediction accuracy by the PRS can vary due to different environmental exposures and gene-environment interactions^19^.

In the present study, we present formulae for estimating polygenic risk score distribution parameters based upon allele frequencies, disease association effect sizes for each SNP, and the SNPxSNP LD matrix. The allele frequencies for different populations can be found in publicly available databases. SNP selection for the PRS can also be informed by the LD structure which is specific to the population of interest via publicly available data (e.g. 1000 genomes). With the caveat that most of the disease risk loci are likely to be shared between populations, our formulae can already be used for assessing disease risk for individuals from a European population and estimate the disease risk for individuals from other populations.

We demonstrate the precision of our formulae with simulations. We show the utility of our estimates in application to AD in the Alzheimer’s Disease Neuroimaging Initiative (ADNI) study^20,21^. We present population specific PRSs for different populations using 1000 Genomes data^22^.

## Material and Methods

### 1000 Genomes project

The aim of this international project is to construct a foundation dataset for human genetics. The data is publicly available at www.1000genomes.org/. It has been used as a main platform that represents shared genetic variation among populations, studying not only common variants that are shared across the world but also rare variation typically restricted to a single continental group^22,23^.

The individuals for the 1000 genomes project were sampled from 19 populations that can be broadly classified as 5 super populations: EUR –European, AMR-ad-mixed Americans, EAS –east Asian, SAS-south Asian, AFR-African (http://www.1000genomes.org/). For our study we used 1583 samples that were genotyped on the Illumina HumanOmni 2.5M BeadChip array. We restricted our analyses to the three largest populations EUR (555 individuals), AFR (457 individuals) and ASN (571 individuals). To make the population-based sample as homogenous as possible, we ran principal component analysis (PCA) using independent SNPs and examined the PCA plot (see Supplementary Figure 1). We excluded 29, 105 and 55 individuals from EUR, AFR, and ASN, respectively, which provided three clear clusters based on the re-derived first two principal components (see Supplementary Figure 2), retaining 526, 352 and 516 individuals, respectively.

SNPs were excluded if their Hardy-Weinberg equilibrium p-value was □10^−6^, the missing value rate was greater than 2% per SNP or the minor allele frequency MAF≤1%, separately in each population.

To choose the set of SNPs mostly associated with AD risk from IGAP summary statistics that is representative in each population, LD pruning was performed in each population separately with parameters r^2^=0.01 and r^2^=0.1 and using a 1000kb window. Then PRSs were generated separately in each population with p-value thresholds p≤10^−5^, 10^−3^, 0.1, 0.5.

### Alzheimer’s Disease Neuroimaging Initiative (ADNI) dataset

ADNI is a public database (http://www.loni.ucla.edu/ADNI/) that was initiated in 2003 by the National Institute of Ageing. The goal of the study is to track the progression of AD using biomarkers, imaging and clinical information along with the genotypic profile. About 900 individuals have different types of longitudinal data; including a clinical diagnosis at each time point of cognitively normal, mild cognitively impaired (MCI) or AD, and an age range between 55-90 years old. 770 samples from ADNI1/GO/2 were whole-genome sequenced (WGS) at high coverage using the Ilumina Omni 2.5M BeadChip (42,732,452 variants). WGS calls were made using the Broad Institute best practices (BWA & GATK HaplotypeCaller). The latest available clinical diagnosis in the ADNI data was used to identify 174 AD cases and 224 cognitively normal individuals for whom genetic information is available. Details of the ADNI study design, participant recruitment, clinical testing, and additional methods have been previously reported elsewhere ^20,21^.

Genetic data was filtered with the same QC steps as for 1000genome data (see above) and matched with the publicly available IGAP GWAS AD summary statistics^24^, retaining 5,771,686 SNPs.

Population structure has been assessed with principal component analysis (PCA) and PCA outliers were excluded, leaving 172 AD cases and 221 cognitively normal individuals for the analyses. PRS were generated with p-value informed clumping (p≤10^−5^, 10^−3^, 0.1, 0.5) with LD pruning parameters r^2^=0.01 and r^2^=0.1 and using a 1000kb window.

### Theoretical estimation of PRS distribution parameters

The PRS approach aggregates the effect of multiple genetic markers identified through Genome-Wide Association Studies (GWAS). For easy interpretation the PRS is often standardised. It is expected that the mean of the PRS in cases will be higher than that in controls, indicating a higher genetic risk for the disorder, but the difference may be small. The standardization requires an estimate of the sample-specific mean and standard deviation (or variance).

If SNPs are independent, we can estimate the mean and the variance of the PRS distribution without knowledge of individual genotypes, simply using the SNP effect sizes and allele frequencies for the SNPs contributing to the PRS, which are often available from the public resources. In the case where SNPs are dependent, estimation of the PRS variance also requires knowledge of the SNPxSNP correlation matrix or a suitable proxy.

Formally, the polygenic risk score for each individual *j*, is

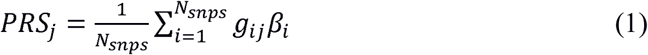

where *j* = 1,…,*N*_*ind*_, *N*_*ind*_ and *N*_*snps*_ are the numbers of individuals and SNPs contributing to the PRS respectively, *g*_*ij*_ is the genotype of SNP *i* for individual *j*, which can take values 0, 1 or 2, and *β*_*i*_ is the effect size of SNP *i* obtained from an independent GWAS study for the disease of interest.

Then the corresponding sample mean and variance can be estimated:

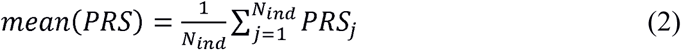

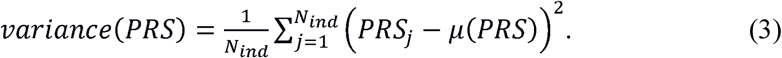

To derive the expected mean and variance of the PRS distribution in terms of effect sizes and SNPs allele frequencies contributing to the PRS, we assume that allele frequencies in a random mating population (Hardy-Weinberg equilibrium) follow a binomial distribution. Then, considering the PRS given as 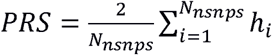. *β*_*i*_ with a Bernoulli-distributed random variable *h*_*i*_ of mean *f*_*i*_, where *f*_*i*_ is the effect allele frequency, the expected mean of the PRS distribution is

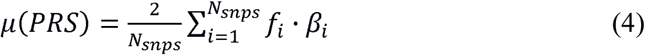

The coefficient “2” in the above formula accounts for the two alleles per SNP. The variance for the product *h*_*i*_ ⋅ *β*_*i*_ for each SNP *i* is 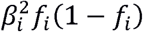, as *f*_*i*_(1 − *f*_*i*_) is the variance of the Bernoulli distributed random variable h_i_. The variance for the total PRS is

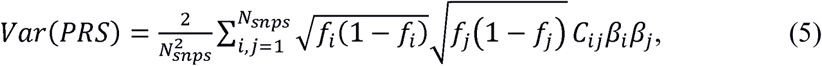

where *C*_*ij*_ is the SNPxSNP correlation between SNP *i* and SNP *j*. If the SNPs are independent, the formula for variance simplifies to

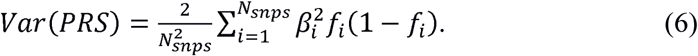

## Results

The standard practice for calculating the PRS is to LD-prune SNPs prior to the analyses. Formulae (4) and (6) offer a very quick estimation of the PRS distribution parameters.

We first examined the performance of the direct calculation of the PRS distribution using formulae (4) and (6) in simulated data. Independent SNPs genotypes were simulated for 1000 individuals with log(OR)~Gaussian(0.01, 0.05) and effect allele frequencies ~Uniform(0.05-0.5). Figure 1 shows PRS density estimated directly from the data (red line) and PRS density calculated using formulae (4), (5) (green line) for a different number of independent SNPs: (a) 100 SNPs, (b) 1000 SNPs and (c) 10000 SNPs.

**Figure 1:**
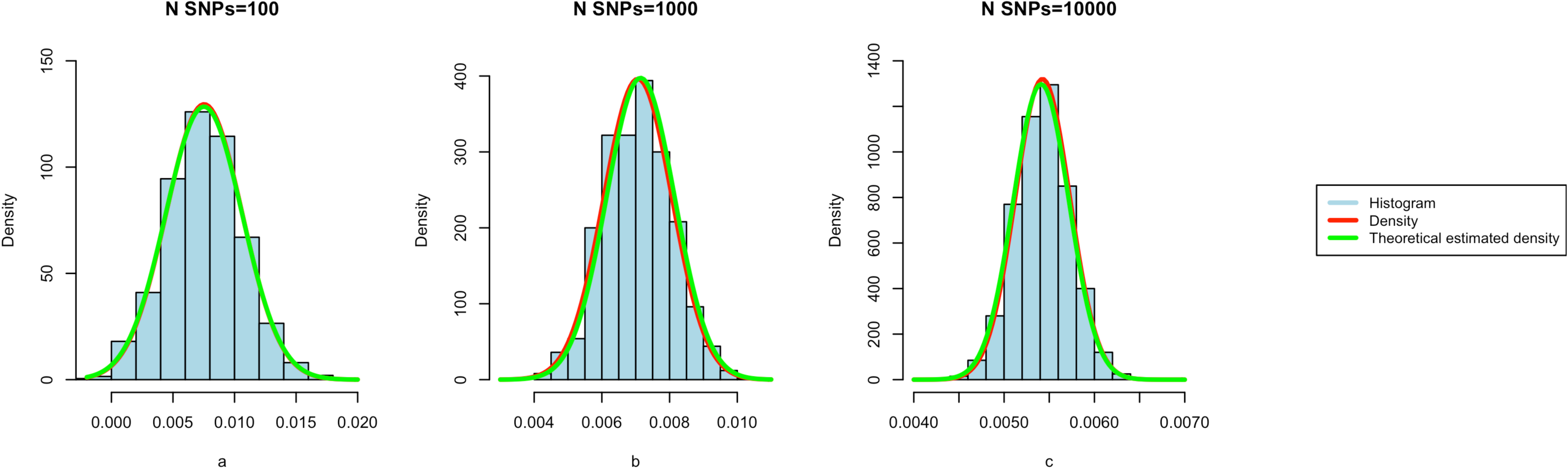
Fitted and estimated PRS densities for simulated genotypes for 1000 individuals. The data was simulated for 1000 individuals with the number of SNPs: (a)- 100, (b)- 1000 and (c)- 10,000. The disease association effect sizes were simulated as log(OR)~N(0.01, 0.05) and allele frequencies were uniformly distributed on the interval [0.05-0.5]. The red curve is the fitted density for the simulated data the green curve is the density with the estimated parameters ((f4), (f6)).

To illustrate the performance of the theoretically estimated PRS parameters with formulae (4) and (6) in real data, we tested it in the 1000 genome dataset for three distinct populations: European, African and Asian. To derive PRS we used publicly available AD GWAS summary statistics^24^ that have been derived in a Caucasian population. The number of LD independent SNPs (r^2^=0.01) contributing to the PRS at different AD significance thresholds in each population, are shown in Table 1. It can be observed that the European population has on average a smaller proportion of significantly associated independent SNPs, as compared to the Chinese and African populations^25^, whereas the total number of SNPs was the largest in the European population. The latter can be explained by the fact that the AD GWAS which was used for the SNP selection procedure is European based and therefore SNP coverage was widest in the European population.

**Table 1.** Number of SNPs in each population (EUR, ASN, AFR) before and after LD pruning. The 1^st^ column shows AD p-value thresholds, the other columns show the number and fraction of SNPs left after pruning for LD with AD GWAS^24^ using parameters r^2^=0.01 and 1Mb window

| **P-value threshold** | **EUR (N=526)**<br>N SNPs<br>independent/total | **ASN (N=516)**<br>N SNPs<br>independent/total | **AFR (352)**<br>N SNPs<br>independent/total |
| --- | --- | --- | --- |
| 1e-5 | 43/346 (12%) | 53/319 (17%) | 56/338 (17%) |
| 1e-3 | 828/2,933 (28%) | 744/2,555 (29%) | 833/2,740 (30%) |
| 0.1 | 15355/146359 (10%) | 14797/122220 (12%) | 13251/135944 (10%) |
| 0.5 | 29839/660922 (5%) | 26973/550861 (5%) | 22320/612208 (4%) |
| 1 | 36618/1452719 (2.5%) | 32055/1266924 (2.5%) | 26329/1633096 (1.6%) |

Figure 2 shows the distributions of non-standardised PRSs, calculated from SNPs with AD association p-value of at least 0.5, in the European, African and Asian populations. It can be seen that the PRS in different populations are not comparable if calculated with standard PRS methods, e.g. the African population can appear to be at high risk for AD since it lies at the very right extreme as compared to the European population. Of course, the differences here are driven only by the genetic background of theses populations, as we used the same summary GWAS statistics for PRS derivation.

**Figure 2:**
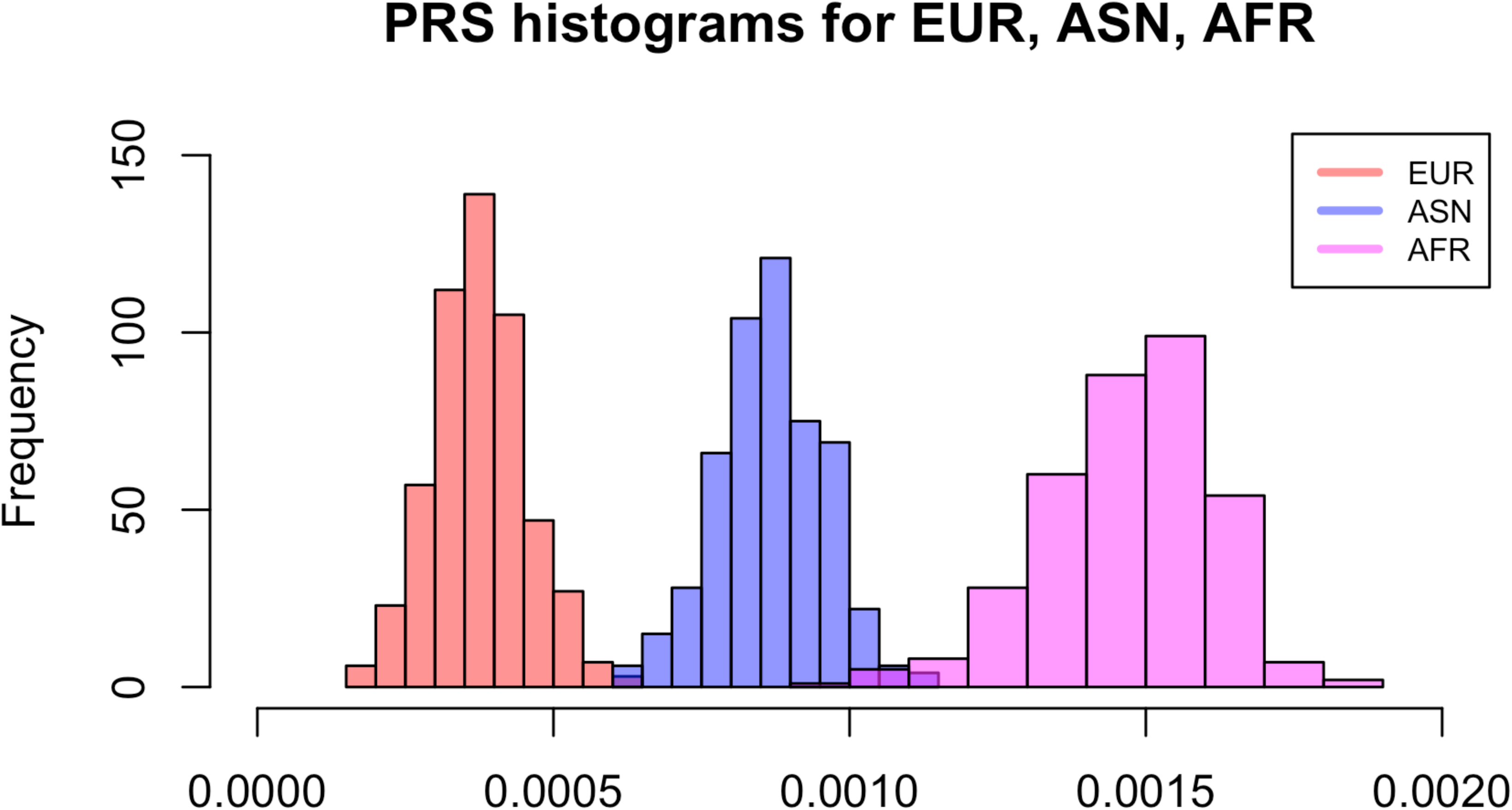
Comparison of PRS distributions in EUR, ASN, AFR populations derived with AD associated SNPs at p-value threshold of 0.5. The PRS were calculated separately for each population (European, Asian and African) with LD pruning parameters r^2^=0.01 and 1Mb window.

The formulae (4) and (6) allow the calculation of the expected mean and variance of the PRS distribution directly for each population, assuming that the SNPs are (almost) independent and SNP effect sizes (i.e., the disease-associated SNP-risks) are similar between populations. The latter presupposes that the biological mechanisms for a particular disease development are generally the same, but we do not exclude that the genome variation and structure vary in different populations. Table 2 displays the PRS distribution parameters (mean and variance) calculated from the sample and estimated using the formulae (4) and (6). The estimates are very similar within the European, Asian and African populations for each of the AD associated p-value thresholds p≤10^−5^, 10^−3^, 0.1, 0.5 and with LD pruning parameter r^2^=0.01. Note that in African population for p≤0.1, 0.5 the variance calculated by formula (6) is underestimated suggesting that some unaccounted SNPxSNP correlation remains.

**Table 2.** The PRS distribution parameter estimates in European, African and Asian populations where the SNPs are LD pruned with r^2^=0.01. The 1^st^ column shows GWAS AD p-value thresholds, the other columns show the number of included SNPs in each population (EUR, ASN and AFR), the PRS mean calculated from the data vs. estimated mean using formulae (f4), and the PRS variance calculated from the data vs. estimated variances using formulae (f5-f6).

| P-value threshold | EUR |  |  | ASN |  |  | AFR |  |  |
| --- | --- | --- | --- | --- | --- | --- | --- | --- | --- |
|  | N SNPs | Actual mean and Estimated mean (f4) | Actual var and Estimated var (f6) var (f5) | N SNPs | Actual mean and Estimated mean (f4) | Actual var and Estimated var (f6) var (f5) | N SNPs | Actual mean and Estimated mean (f4) | Actual var and Estimated var (f6) var (f5) |
| 1e-5 | 43 | 9.4e-3<br>9.3e-3 | 8.9e-5<br>7.6e-5<br>7.6e-5 | 53 | 8.4e-3<br>8.5e-3 | 6.4e-5<br>6.1e-5<br>6.1e-5 | 56 | 4.9e-3/<br>5e-3 | 7.7e-5<br>8.2e-5<br>8.3e-5 |
| 1e-3 | 828 | 1.2e-3<br>1.2e-3 | 8.8e-7<br>8.1e-7<br>8.3e-7 | 744 | 2.4e-3<br>2.5e-3 | 1.16e-6<br>1.12e-6<br>1.15e-6 | 833 | 4.9e3/<br>4.9e-3 | 1.3e-6<br>1.2e-6<br>1.23e-6 |
| 0.1 | 15355 | 6.2e-4<br>6.2e-4 | 1.86e-8<br>1.63e-8<br>2.3e-8 | 14797 | 1.3e-3<br>1.3e-3 | 2.3e-8<br>2.1e-8<br>2.9e-8 | 13251 | 2e-3/<br>2e-3 | 4.5e-8<br>2.7e-8<br>4e-8 |
| 0.5 | 29839 | 3.7e-4<br>3.7e-4 | 6.3e-9<br>5.4e-9<br>8.2e-9 | 26973 | 8.6e-4<br>8.6e-4 | 8.2e-9<br>7.4e-9<br>1e-8 | 22320 | 1.5e-3/<br>1.5e-3 | 1.9e-8<br>1.1e-8<br>1.8e-8 |

When the SNPs are in noticeable LD, the estimation of the PRS variance (5) requires knowledge of the SNPxSNP correlation matrix, which in turn requires the availability of individual genotypes. This complicates the application of formula (5) to real data, as the computational burden for estimation of the full correlation matrix may be substantial.

In practice, LD-intelligent pruning is performed before PRS calculation to pick up disease-relevant independent variants. Typically the threshold squared correlation parameter (r^2^) varies from 0.1 to 0.2 in a 500kb or 1000kb window. Of course, this procedure does not make SNPs absolutely independent and some correlation remains. If this correlation is ignored, then the PRS variance is underestimated. We have compared the variance estimates with our formulae for threshold values r^2^=0.01 (r=0.1) and r^2^=0.1 (r=0.316). As expected, the most stringent pruning (r^2^=0.01) provides a closer resemblance of the theoretically estimated variance (by formula (6)) to the estimate assuming dependence (by formula (5)).

When SNPs are pruned for LD in a window, say 1000kb, only a small proportion of SNPs remains correlated in that window. For example, out of ~10M SNPs in 1000Genomes European data, ~100K SNPs remained after r^2^=0.01 pruning and ~900K SNPs remained after r^2^=0.1 pruning. In a window of 1MB, there were on average 3,000 SNPs before pruning, and ~300 SNPs after r^2^=0.1 pruning. About 30% of SNP pairs had correlation less than r^2^=0.0001, ~93% of SNP pairs had LD less than r^2^=0.01, and only 1% had r^2^ greater than 0.1, the pruning threshold. Thus, very roughly, the average correlation per LD block is between 0.002 and 0.003.

Applying r=0.002 as an approximation of correlation between SNPs in formula (5) provides closer estimates of PRS variance to the data-derived variance for all three populations in 1000 genomes project including the African population, see Table 2, the variance estimated by formula (5). Note that this coefficient works also well for other pruning parameters of r^2^=0.01, 0.1 (see Tables 2,3 and Figures 3,4).

**Table 3.**
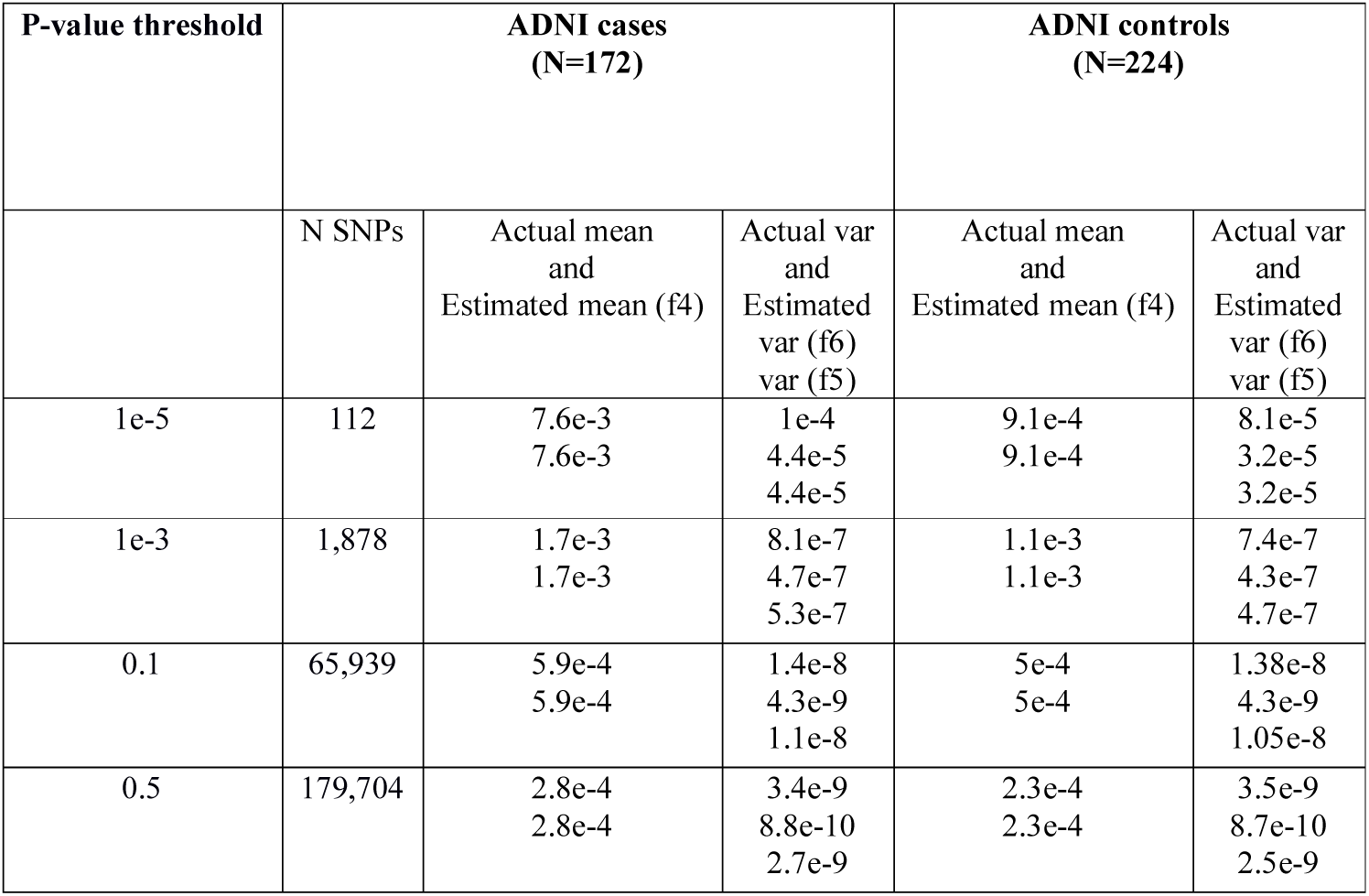
The PRS distribution parameter estimates (formulae (f4-f6)) in ADNI dataset for AD cases and controls with LD pruned SNPs (r^2^=0.1). The 1^st^ column shows GWAS AD p-value thresholds, the other columns show the PRS mean calculated from the data vs. estimated mean using formulae (f4), and the PRS variance calculated from the data vs. estimated variances using formulae (f5-f6).

**Figure 3:**
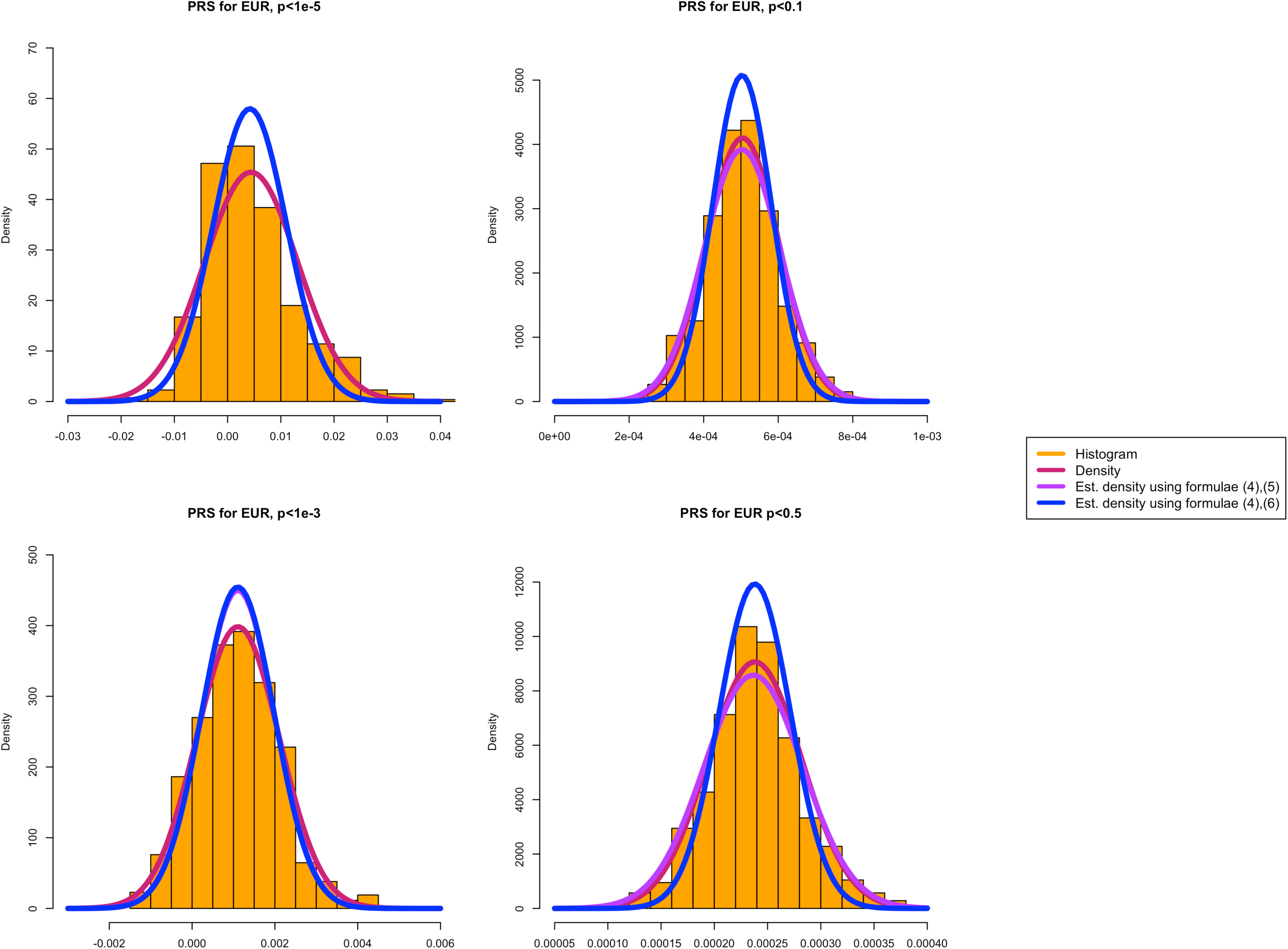
Sample-based and estimated (formulae (4), (5-6)) PRS densities in 1000 genomes for different AD association p-value thresholds of 10^−5^, 10^−3^, 0.1, 0.5. A: European population B: Asian population C: African population

**Figure 4:**
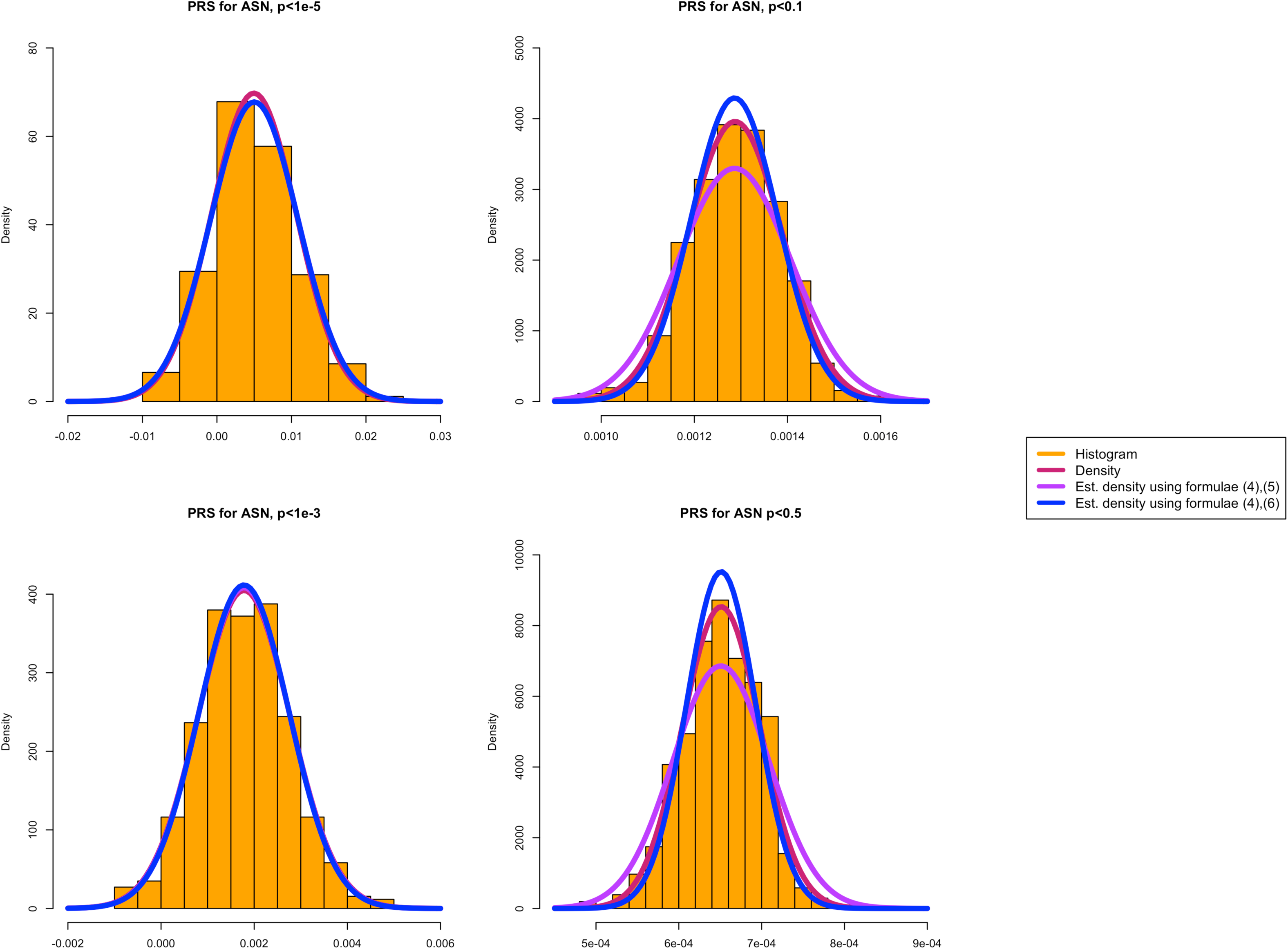
Sample-based and estimated PRS densities in AD cases and controls of ADNI dataset with LD pruning parameters r^2^=0.01 and 1Mb window for different AD association p-value thresholds of 10^−5^, 10^−3^, 0.1, 0.5.

In order to demonstrate the accuracy of the proposed approximation, we use the LD pruning parameter of r^2^=0.1, which is commonly used in practice. Figure 3: a) - European, b)-Asian and c)-African populations shows the fitted distributions for all three methods: actual PRS densities, PRS density estimated by formulae (4),(6) and PRS density estimated by formulae (4),(5) where SNPxSNP correlation matrix is estimated by r=0.002. Remarkably, even with pruning threshold r^2^=0.1 the variance is severely underestimated when LD is neglected (blue curves); the estimated parameters provide much better fit to the real data when LD is taken into account (purple curves).

Finally, we demonstrated the performance of our theoretical formulae in AD case/control data (ADNI). First, we used a quite stringent LD pruning parameter r^2^=0.01 and calculated the PRS distribution parameters separately for 172 AD cases and 221 cognitively normal controls. Figure 4 shows histograms of the PRSs and their density curves as compared to the PRS densities whose parameters are calculated with formulae (4), (5-6). A better fit can be observed for all p-value thresholds with the formula (5) which accounts for potentially remaining LD between SNPs. Table 3 presents the results when the LD pruning parameter r^2^ was set to r^2^=0.1. In this case, accounting for the LD (i.e. using formula (5)) is crucial.

## Discussion

In this study we have derived theoretical formulae to estimate the parameters of any polygenic risk score distribution. The proposed estimations depend on the allele frequencies and the effect sizes of the risk alleles for a particular disorder, both are often publicly available. The presented formula (6) assumes that the SNPs are independent, and the allele frequencies are derived from a homogenous population. If these assumptions are satisfied, our estimates are quite accurate for both simulated and real data. The SNPs which are used for the generation of PRSs are typically LD pruned, which is specific to each population LD structure. There are publicly available datasets (e.g. 1000 genomes), where the pruning can be performed with respect to the population and disease of interest. If the SNPs contributing to the PRS are not LD independent, then the variance of the PRS distribution is underestimated. In reality even after a stringent LD pruning, some correlation between SNPs remains. We recommend using formula (5) for PRS variance estimation to account for remaining LD in all cases.

We verified our formulae using simulations and then applied it to the 1000 genomes data and the ADNI dataset. We showed that the PRSs for AD differ substantially in different populations even assuming that the risk loci are the same. For a given set of disease association effect sizes, our formulae provide extremely close estimates for the PRS distribution parameters without calculating individual scores in a sample in simulated and real data from European, African and Chinese populations from 1000 genomes data. It also shows very good approximation for AD cases and cognitively normal individuals controls in the ADNI dataset.

The ability to estimate the expected parameters of the PRS distribution in a population without access to the protected individual genotypes of cases and controls have great potential in practice. Firstly, it can be used to add controls to studies without requiring genotyping, if population-based controls are suitable (for example when the prevalence of the disease in the population is not too high). Secondly, often the raw genotypes cannot be shared between collaborators, but the individual polygenic risk scores and the SNPs which were used to derive them, are easier to share. The proposed theoretical estimates can be used for harmonisation of PRSs derived in studies where the data were generated with different chips and/or the QC steps were different, resulting in PRSs calculated from different SNP sets. Thirdly, for risk prediction tools/software, algorithms based on our formulae can compare an individual’s score (for the whole genome and/or for a specific gene set) against the expected PRS distribution and thus potentially predict the disease risk of a single individual without accessing a large case/control background sample. Fourthly, if we assume that the disease risk effect sizes are similar in different populations, it is possible to predict the disease risk in a population using the effect sizes from Caucasian studies. Of course, this approach is not perfect as the effect sizes may also differ between populations, however, it can now inform non-Caucasian study designs in the absence of information as at least two (allele frequencies and LD structure) out of three unknowns in the equation are taken care of. In addition, if correction for LD is required, the SNPxSNP LD matrix needs to be estimated, which can be computationally demanding. In the Results section we suggested some approximation of LD for commonly used pruning parameters, but this LD approximation will differ depending on the sets of SNPs of interest to the researchers (e.g. whole genome or set of genes in a pathway, either imputed or genotyped, etc.). And finally, these formulae are very simple, and the computation time is negligible, which allows researchers to make quick decisions about the suitability of their in-house samples for their studies without lengthy procedures for external data application and processing. For example, if a research team is interested in selecting cell lines which are enriched for neuroimmunology related biological pathways, they can generate PRS for their existing cell lines and compare the derived pathway-specific polygenic scores against expected PRS mean in the population for this particular pathway.

## Contributions

GL, EG, KMS, VEP wrote the manuscript and prepared figures and tables. All authors reviewed and approved the final version of the manuscript.

## Acknowledgements

We thank the MRC Centre for Neuropsychiatric Genetics and Genomics and UK Dementia Research Institute (DRI), Dementia Platform UK (DPUK).

Also we thank ADNI database. Data collection and sharing for this project was funded by the Alzheimer’s Disease Neuroimaging Initiative (ADNI) (National Institutes of Health Grant U01 AG024904) and DOD ADNI (Department of Defense award number W81XWH-12-2-0012). ADNI is funded by the National Institute on Aging, the National Institute of Biomedical Imaging and Bioengineering, and through generous contributions from the following: AbbVie, Alzheimer’s Association; Alzheimer’s Drug Discovery Foundation; Araclon Biotech; BioClinica, Inc.; Biogen; Bristol-Myers Squibb Company; CereSpir, Inc.; Cogstate; Eisai Inc.; Elan Pharmaceuticals, Inc.; Eli Lilly and Company; EuroImmun; F. Hoffmann-La Roche Ltd and its affiliated company Genentech, Inc.; Fujirebio; GE Healthcare; IXICO Ltd.; Janssen Alzheimer Immunotherapy Research & Development, LLC.; Johnson & Johnson Pharmaceutical Research & Development LLC.; Lumosity; Lundbeck; Merck & Co., Inc.; Meso Scale Diagnostics, LLC.; NeuroRx Research; Neurotrack Technologies; Novartis Pharmaceuticals Corporation; Pfizer Inc.; Piramal Imaging; Servier; Takeda Pharmaceutical Company; and Transition Therapeutics. The Canadian Institutes of Health Research is providing funds to support ADNI clinical sites in Canada. Private sector contributions are facilitated by the Foundation for the National Institutes of Health (www.fnih.org). The grantee organization is the Northern California Institute for Research and Education, and the study is coordinated by the Alzheimer’s Therapeutic Research Institute at the University of Southern California. ADNI data are disseminated by the Laboratory for Neuro Imaging at the University of Southern California.

Acknowledgements to funders: DRI, DPUK, MRC Centre for Neuropsychiatric Genetics and Genomics

---

**Supplementary Figure 1.**
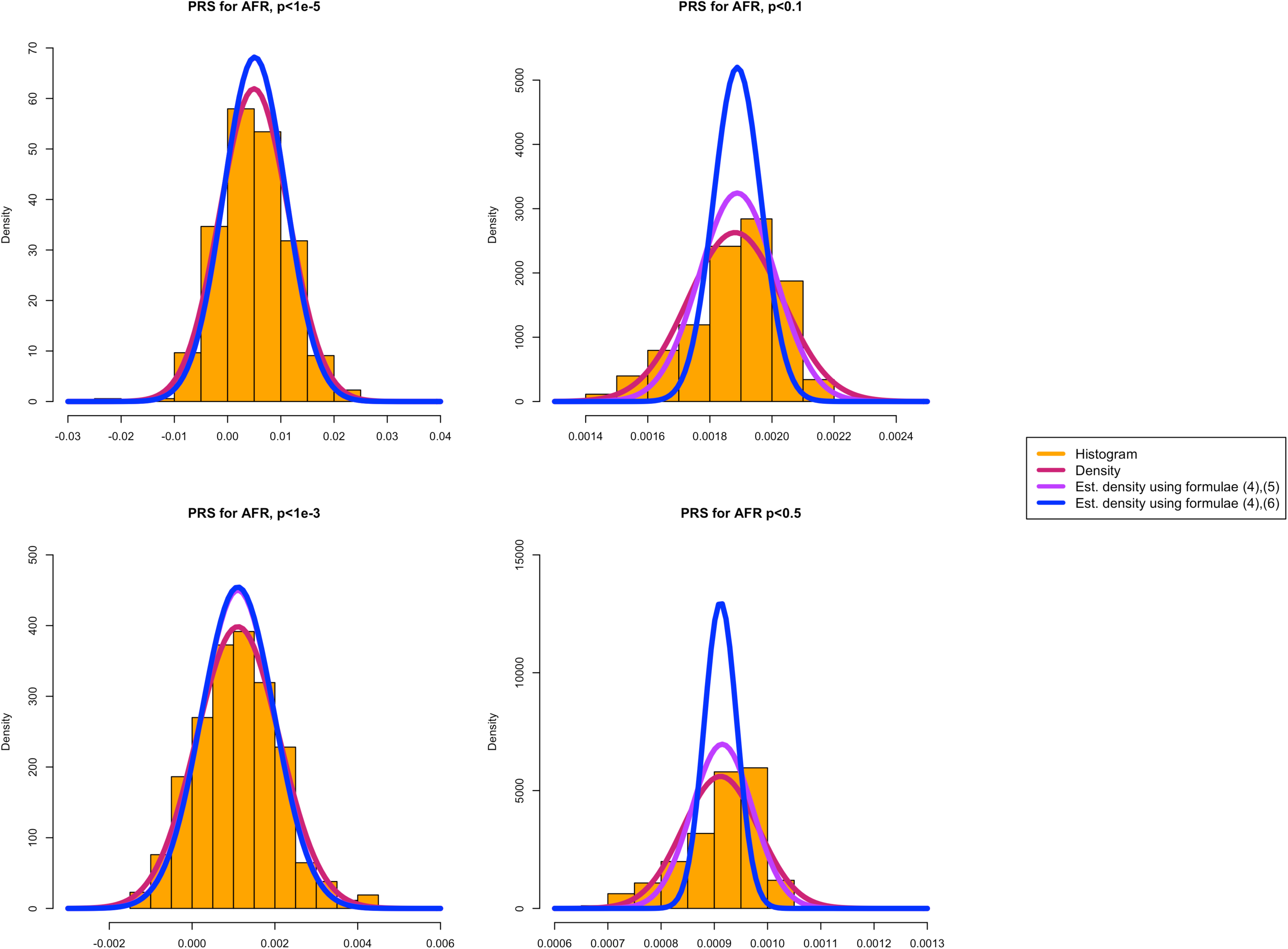
First two PCA components in not QCed 1000genome dataset for the three ancestry groups. First two PCA are plotted for individuals from 1000genome dataset for the three ancestry groups: 555-European, 457-African and 571-Asian.

**Supplementary Figure 2.**
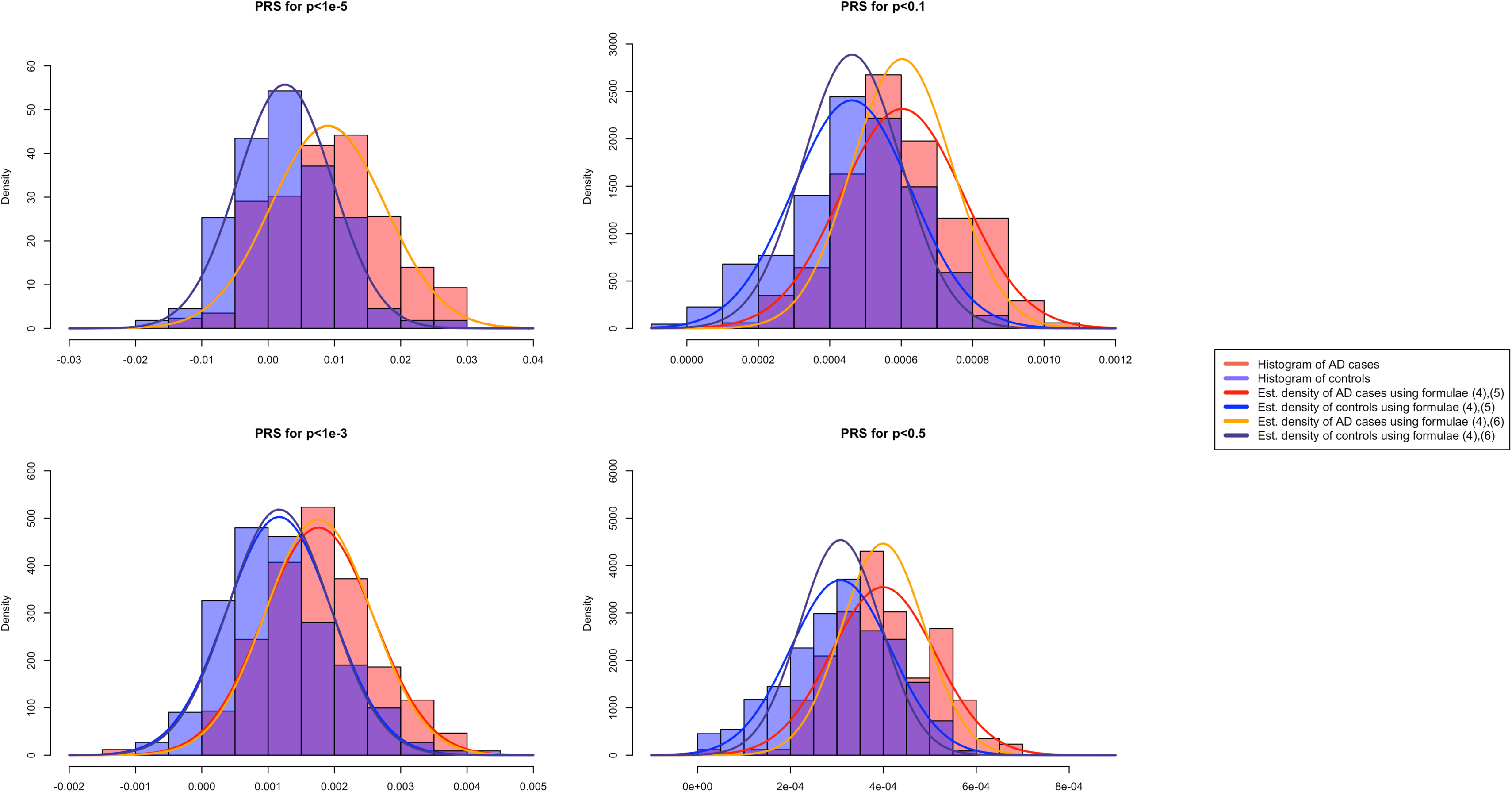
First two PCA components in QCed 1000genome dataset for the three ancestry groups. First two PCA components are plotted for individuals (after removing PCA outliers) from 1000genome dataset for the three ancestry groups: 526-European, 352-African and 516-Asian.

