## Supplementary figures and images for "Positioning Personal Polygenic Risk score against the population background"

### Supplementary Figure 1

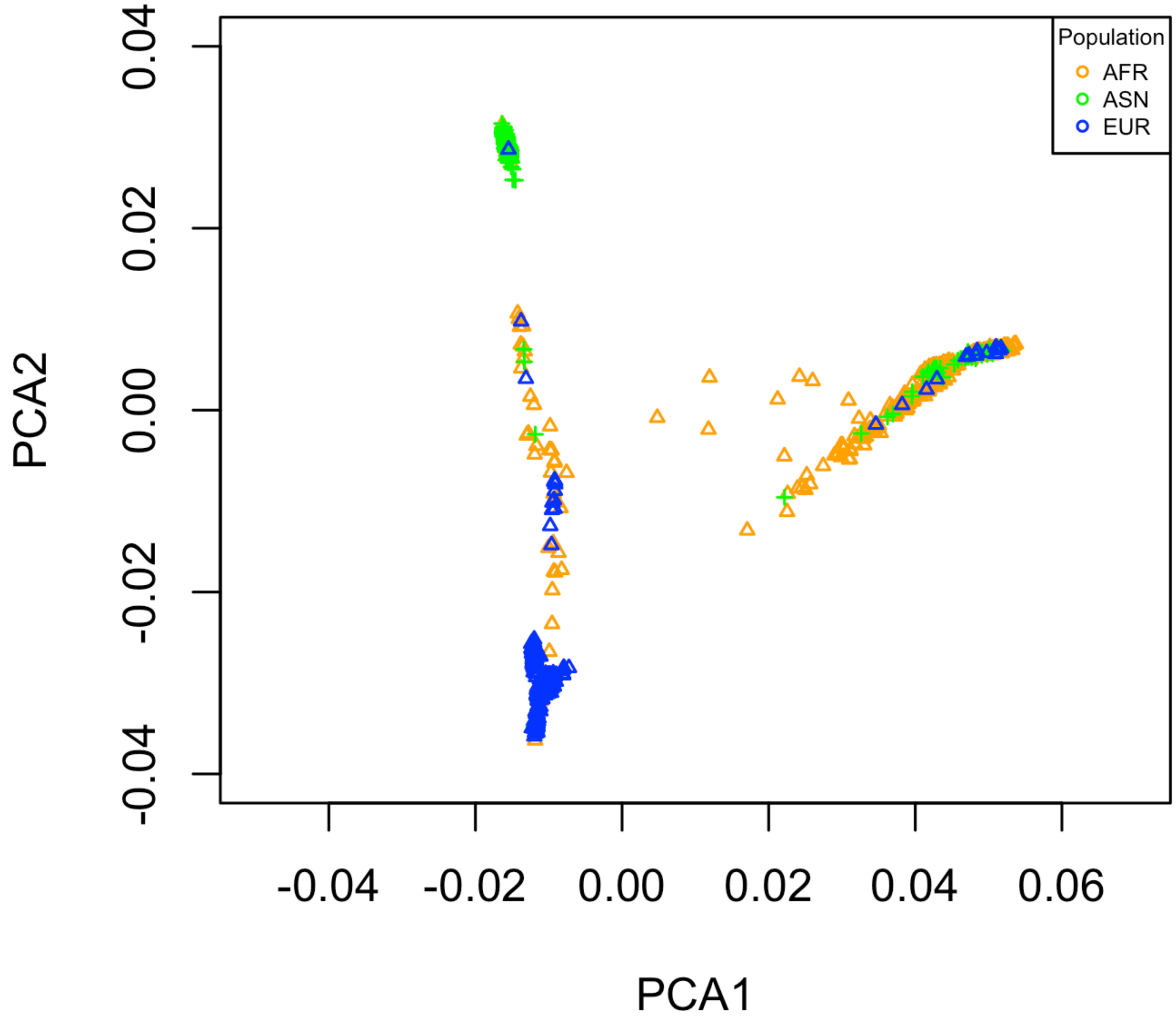

### Supplementary Figure 2

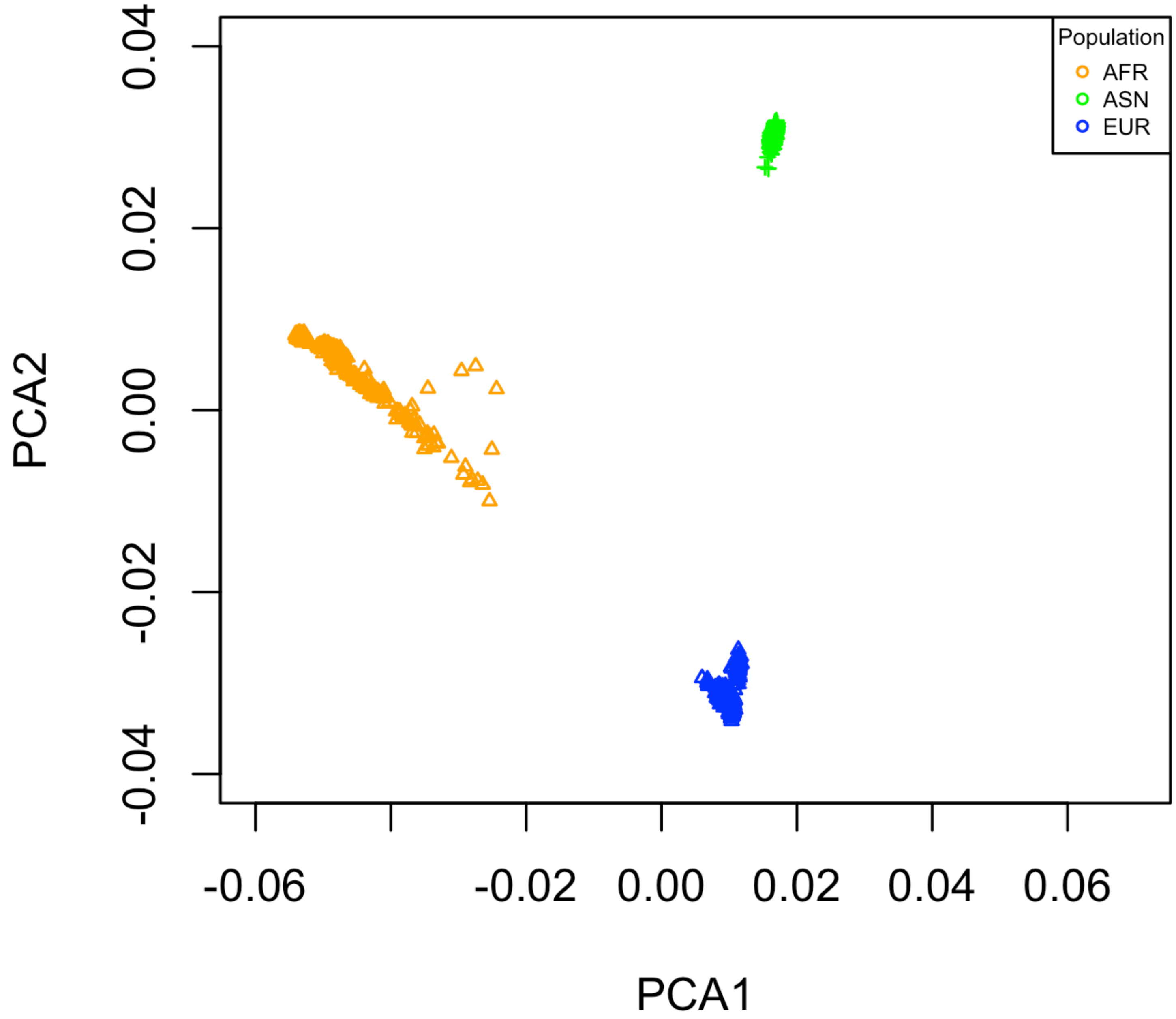
